# A novel toxin-antitoxin module SlvT–SlvA governs megaplasmid stability and incites solvent tolerance in *Pseudomonas putida* S12

**DOI:** 10.1101/2020.01.02.893495

**Authors:** Hadiastri Kusumawardhani, David van Dijk, Rohola Hosseini, Johannes H. de Winde

## Abstract

*Pseudomonas putida* S12 is highly tolerant towards organic solvents in saturating concentrations, rendering this microorganism suitable for the industrial production of various aromatic compounds. Previous studies reveal that *P. putida* S12 contains a single-copy 583 kbp megaplasmid pTTS12. This pTTS12 encodes several important operons and gene clusters facilitating *P. putida* S12 to survive and grow in the presence of toxic compounds or other environmental stresses. We wished to revisit and further scrutinize the role of pTTS12 in conferring solvent tolerance. To this end, we cured the megaplasmid from *P. putida* S12 and conclusively confirmed that the SrpABC efflux pump is the major contributor of solvent tolerance on the megaplasmid pTTS12. Importantly, we identified a novel toxin-antitoxin module (proposed gene names *slvT* and *slvA* respectively) encoded on pTTS12 which contributes to the solvent tolerant phenotype and is essential in conferring genetic stability to the megaplasmid. Chromosomal introduction of the *srp* operon in combination with *slv*AT gene pair created a solvent tolerance phenotype in non-solvent tolerant strains such as *P. putida* KT2440, *E. coli* TG1, and *E. coli* BL21(DE3).

**Importance:** Sustainable alternatives for high-value chemicals can be achieved by using renewable feedstocks in bacterial biocatalysis. However, during bioproduction of such chemicals and biopolymers, aromatic compounds that function as products, substrates or intermediates in the production process may exert toxicity to microbial host cells and limit the production yield. Therefore, solvent-tolerance is a highly preferable trait for microbial hosts in the biobased production of aromatic chemicals and biopolymers. In this study, we revisit the essential role of megaplasmid pTTS12 from solvent-tolerant *P. putida* S12 for molecular adaptation to organic solvent. In addition to the RND efflux pump (SrpABC), we identified a novel toxin-antitoxin module (SlvAT) which contributes to tolerance in low solvent concentration as well as to genetic stability of pTTS12. These two gene clusters were successfully transferred to non-solvent tolerant strains of *P. putida* and to *E. coli* strains to confer and enhance solvent tolerance.

## Introduction

One of the main problems in the production of aromatic compounds is chemical stress caused by the added substrates, pathway intermediates, or products. These chemicals, often exhibiting characteristics of organic solvents, are toxic to microbial hosts and may negatively impact product yields. They adhere to the cell membranes, alter membrane permeability, and cause membrane damage (1, 2). *Pseudomonas putida* S12 exhibits exceptional solvent tolerance characteristics, enabling this strain to withstand toxic organic solvents in saturating concentrations (3, 4). Consequently, a growing list of valuable compounds has successfully been produced using *P. putida* S12 as a biocatalyst by exploiting its solvent tolerance (5–9).

Following the completion of its full genome sequence and subsequent transcriptome and proteome analyses, several genes have been identified that may play important roles in controlling and maintaining solvent tolerance of *P. putida* S12 (10–12). As previously reported, an important solvent tolerance trait of *P. putida* S12 is conferred through the RND-family efflux pump, SrpABC, which actively removes organic solvent molecules from the cells (13, 14). Initial attempts to heterologously express the SrpABC efflux pump in *E. coli* enabled instigation of solvent-tolerance and production of 1-naphtol (15, 16). Importantly, the SrpABC efflux pump is encoded on the megaplasmid pTTS12 of *P. putida* S12 (12).

The 583 kbp megaplasmid pTTS12 is a stable single-copy plasmid specific to *P. putida* S12 (12). It encodes several important operons and gene clusters enabling *P. putida* S12 to tolerate, resist and survive the presence of various toxic compounds or otherwise harsh environmental conditions. Interesting examples are the presence of a complete styrene degradation pathway gene cluster, the RND efflux pump specialized for organic solvents (SrpABC) and several gene clusters conferring heavy metal resistance (12, 17, 18). In addition, through analysis using TADB2.0 (19, 20) pTTS12 is predicted to contain three toxin-antitoxin modules. Toxin-antitoxin modules recently have been recognized as important determinants of resistance towards various stress conditions (21, 22). Toxin-antitoxin modules identified in pTTS12 consist of an uncharacterized RPPX_26255 -RPPX_26260 system and two identical copies of a VapBC system (23). RPPX_26255 and RPPX_26260 belong to a newly characterized type II toxin-antitoxin pair COG5654-COG5642. While toxin-antitoxin systems are known to preserve plasmid stability through post-segregational killing of plasmid-free daughter cells (24), RPPX_26255-RPPX_26260 was also previously shown to be upregulated during organic solvent exposure indicating its role in solvent tolerance (11).

In this paper, we further address the role of pTTS12 in conferring solvent tolerance of *P. putida* S12. Curing pTTS12 from its host strain caused a significant reduction in solvent tolerance, while complementation of the cured strain with the *srp* operon significantly restored solvent tolerance, underscoring the importance of the SrpABC solvent pump in conferring solvent tolerance in *P. putida* S12. In addition, we showed that the novel toxin-antitoxin pair *slvAT* (RPPX_26260 and RPPX_26255) is essential for maintaining genetic stability of megaplasmid pTSS12. We further modelled SlvT and SlvA to the recently characterised crystal structure of McbT and McbA from *Mycobacterium tuberculosis* (25). We clearly show that SlvT causes toxicity by degrading NAD^+^ in *E. coli* BL21 (DE3), similar to recently characterized toxins of the COG5654 family (25–27). Importantly, introduction of the *srp* operon in combination with toxin-antitoxin pair *slvAT* in non-solvent tolerant *P. putida* KT2440 as well as *E. coli* confers and enhances solvent tolerance in these strains.

## Results

### Megaplasmid pTTS12 is essential for solvent tolerance in *P. putida* S12

To further analyze the role of the megaplasmid of *P. putida* S12 in solvent tolerance, pTTS12 was removed from *P. putida* S12 using mitomycin C. This method was selected due to its reported effectivity in removing plasmids from *Pseudomonas sp.* (28), although previous attempts regarded plasmids that were significantly smaller in size than pTTS12 (29). After treatment with mitomycin C (10-50 mg L^-1^), liquid cultures were plated on M9 minimal media supplemented with indole to select for plasmid-cured colonies. Megaplasmid pTTS12 encodes two key enzymes: Styrene monooxygenase (SMO) and Styrene oxide isomerase (SOI), that are responsible for the formation of indigo coloration from indole. This conversion results in indigo coloration in spot assays for wildtype *P. putida* S12 whereas white colonies are formed in the absence of megaplasmid pTTS12. With the removal of pTTS12, loss of indigo coloration and hence, of indigo conversion was observed in all three plasmid-cured strains and the negative control *P. putida* KT2440 (Figure 1A).

**Figure 1.**
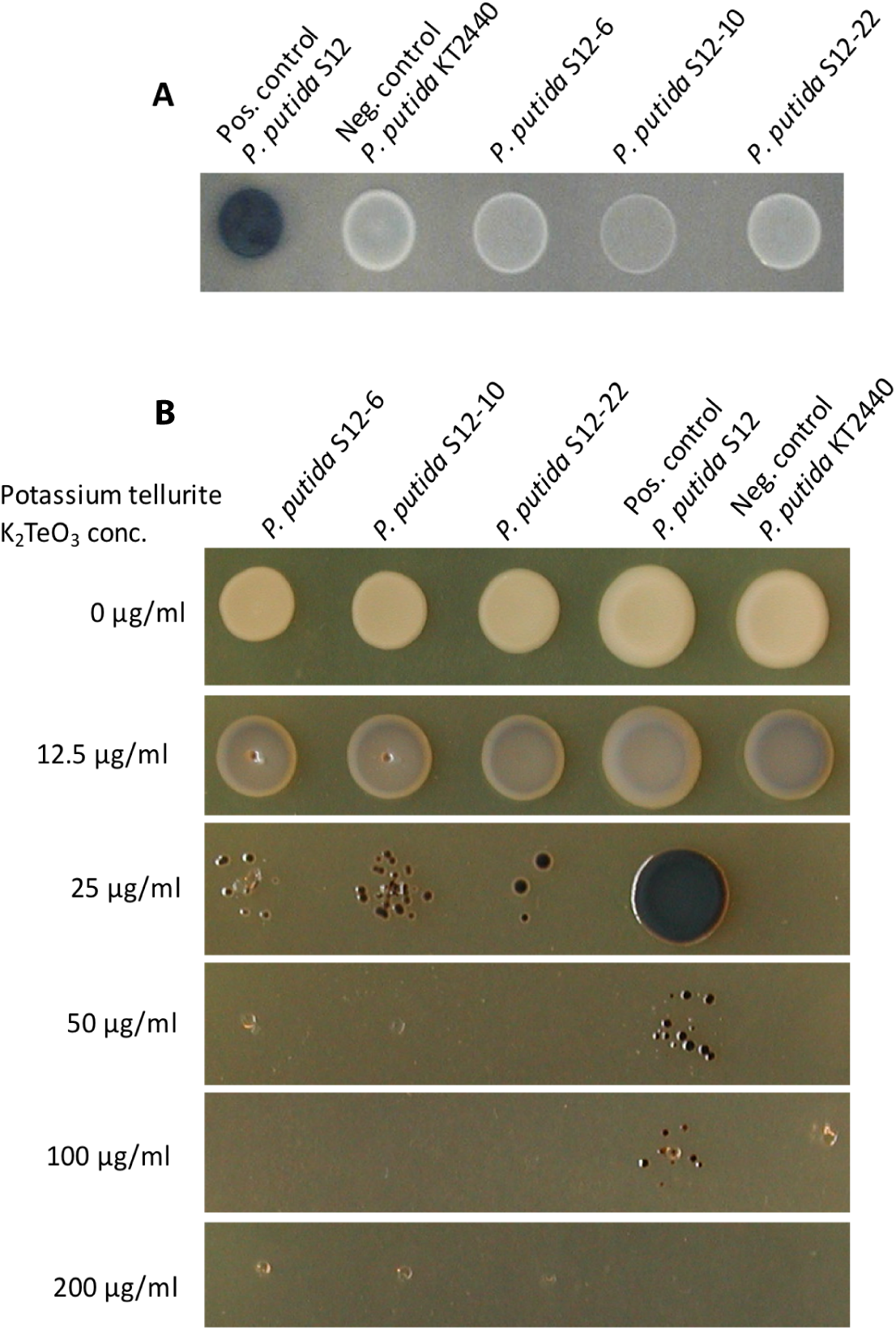
**Curing of the megaplasmid pTTS12 from *P. putida* S12.** A. Activity of styrene monooxygenase (SMO) and styrene oxide isomerase (SOI) for indigo formation from indole in *P. putida* strains. Enzyme activity was lost in the megaplasmid-cured strains S12 ΔpTTS12 (white colonies). Indole (100 mg L^-1^) was supplemented in M9 minimum media. B. K_2_TeO_3_ resistance of *P. putida* strains on lysogeny broth (LB) agar. Tellurite resistance was reduced in the megaplasmid-cured strains S12 ΔpTTS12 (MIC 50 mg L^-1^).

With mitomycin C concentration of 30 mg L^-1^, 3 out of 122 obtained colonies appeared to be completely cured from the megaplasmid, underscoring the high genetic stability of the plasmid. No colonies survived the addition of 40 and 50 mg L^-1^ of mitomycin C, whereas all the colonies that survived the addition of 10 and 20 mg L^-1^ of mitomycin C retained the megaplasmid. All three independent colonies cured from the megaplasmid were isolated as *P. putida* S12-6, *P. putida* S12-10, and *P. putida* S12-22. Complete loss of the megaplasmid was further confirmed by phenotypic analysis (Figure 1), and by full genome sequencing. Several operons involved in heavy metal resistance were previously reported in the pTTS12 (12). The *terZABCD* operon contributes to tellurite resistance in wildtype *P. putida* S12 with minimum inhibitory concentration (MIC) as high as 200 mg L^-1^ (Figure 1B). In the megaplasmid-cured strains, severe reduction of tellurite resistance was observed, decreasing the potassium tellurite MIC to 50 mg L^-1^ (Figure 1B).

Genomic DNA sequencing confirmed complete loss of pTTS12 from *P. putida* strains S12-6, S12-10, and S12-22 without any plasmid-derived fragment putatively being inserted within the chromosome. Complementation of pTTS12 into the plasmid-cured *P. putida* S12 strains restored the indole-indigo transformation and high tellurite resistance to the similar level with wildtype strain (Figure S1). Repeated megaplasmid curing experiments indicated that *P. putida* S12 can survive the addition of 30 mg L^-1^ Mitomycin C with the frequency of 2.48 (± 0.58) x 10^-8^. Among these survivors, only 2% colony population lost the megaplasmid, confirming the genetic stability of pTSS12. In addition, other plasmid-curing attempt by introducing double strand break as described by Wynands and colleagues (30) was not successful due to the pTTS12 stability.

Growth comparison in solid and liquid culture in the presence of toluene was performed to analyze the effect of megaplasmid curing in constituting solvent tolerance trait of *P. putida* S12. In contrast with wildtype *P. putida* S12, the plasmid-cured strains were unable to grow under toluene atmosphere. In liquid LB medium, plasmid-cured *P. putida* S12 strains were able to tolerate a maximum of 0.15% v/v toluene, whereas the wildtype *P. putida* S12 can grow in the presence of 0.30% v/v toluene (Figure 2). In the megaplasmid-complemented *P. putida* S12-C strains, solvent tolerance was restored to the wildtype level (Figure S1-D). Hence, absence of megaplasmid pTTS12 caused a significant reduction of solvent tolerance in *P. putida* S12. We chose *P. putida* S12-6 for further experiments representing megaplasmid-cured *P. putida* S12.

**Figure 2.**
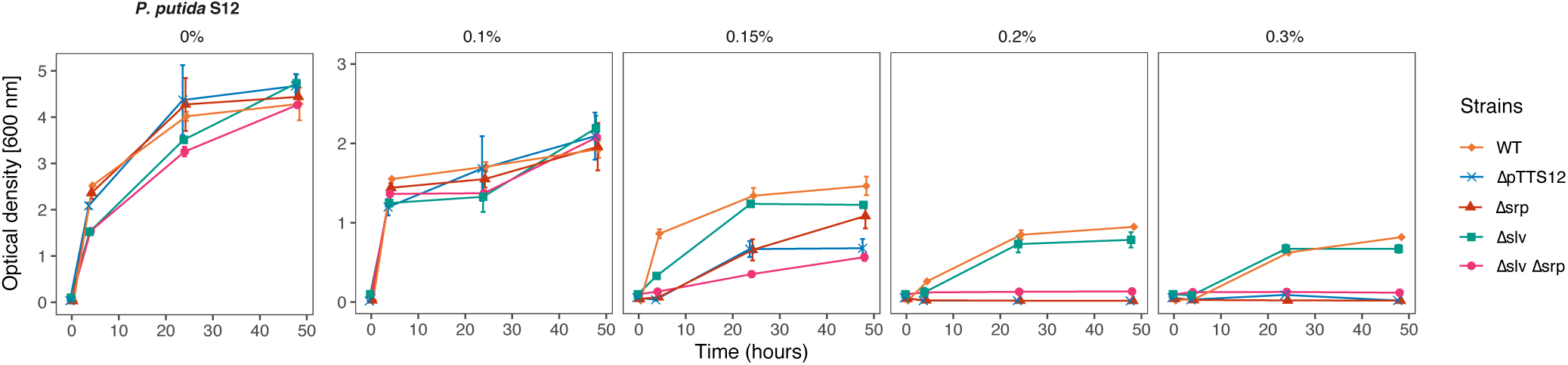
Megaplasmid pTTS12 determines the solvent tolerance trait of *P. putida* S12. Solvent tolerance analysis was performed on wildtype *P. putida* S12, *P. putida* S12 ΔpTTS12, *P. putida* S12 Δsrp, *P. putida* S12 Δslv, and *P. putida* S12 Δsrp Δslv growing in liquid LB media with 0, 0.10, 0.15, 0.20 and 0.30 % v/v toluene. The removal of the megaplasmid pTTS12 clearly caused a significant reduction in the solvent tolerance of *P. putida* S12 ΔpTTS12. Deletion of srpABC (Δsrp), RPPX_26255-26260 (Δslv), and the combination of these gene clusters (Δsrp Δslv) resulted in a lower solvent tolerance. This figure displays the mean of three biological replicates and error bars indicate standard deviation. The range of y-axis is different in the first panel (0–5) than the rest of the panels (0–3).

### The SrpABC efflux pump and gene pair RPPX_26255-26260 are the main constituents of solvent tolerance encoded on pTTS12

The significant reduction of solvent-tolerance in plasmid-cured *P. putida* S12 underscored the important role of megaplasmid pTTS12 in solvent-tolerance. Besides encoding the efflux pump SrpABC enabling efficient intermembrane solvent removal (12, 13), pTTS12 encodes more than 600 genes and hence, may contain multiple additionally solvent-tolerance related genes. Two adjacent hypothetical proteins, RPPX_26255 and RPPX_26260, encoded on the megaplasmid pTTS12 were previously reported to be upregulated in the presence of toluene (11). We propose to name RPPX_26255-26260 gene pair as ‘*slv’* due to its elevated expression in the presence of solvent. In a first attempt to identify additional potential solvent tolerance regions of pTTS12, we deleted the *srp*ABC genes (Δ*srp*), RPPX_26255-26260 genes (Δ*slv*), and the combination of both gene clusters (Δ*srp* Δ*slv*) from pTTS12 in wild-type *P. putida* S12.

All strains were compared for growth under increasing toluene concentrations in liquid LB medium (Figure 2). In the presence of low concentrations of toluene (0.1% v/v), all strains showed similar growth. With the addition of 0.15% v/v toluene, S12 Δ*slv*, S12 Δ*srp* and S12 Δ*srp* Δ*slv* exhibit slower growth and reached a lower OD_600nm_ compared to the wildtype S12 strain. S12 Δ*slv* and S12 Δ*srp* achieved a higher OD_600nm_ in batch growth compared to S12 ΔpTTS12 and S12 Δ*srp* Δ*slv* due to the presence of SrpABC efflux pump or RPPX_26255-26260 gene pair. Interestingly, S12 Δ*srp* Δ*slv* (still containing pTSS12) exhibit diminished growth compared to S12 ΔpTTS12. This may be an indication of megaplasmid burden in the absence of essential genes for solvent tolerance. With 0.2% and 0.3% v/v toluene added to the medium, S12 Δ*srp,* S12 Δ*srp* Δ*slv*, and S12 ΔpTTS12 were unable to grow while the wildtype S12 and S12 Δ*slv* were able to grow although S12 Δ*slv* reached a clearly lower OD_600nm_ compared to wildtype S12. Taken together, these results demonstrate an important role for both the SrpABC efflux pump and the *slv* gene pair in conferring solvent tolerance.

### Transferability of solvent tolerance exerted by SrpABC efflux pump and *slv* gene pair in Gram-negative bacteria

Functionality of the *srp* operon and *slv* gene pair was explored in the model Gram-negative non-solvent tolerant strains, *P. putida* KT2440, *E. coli* TG1 and *E. coli* BL21 (DE3). We complemented *srpRSABC* (*srp* operon), *slv* gene pair, and a combination of both gene clusters into *P. putida* S12-6, *P. putida* KT2440, *E. coli* TG1, and *E. coli* BL21 (DE3) using mini-Tn7 transposition.

Chromosomal introduction of *slv* into S12-6 and KT2440, improved growth of the resulting strains at 0.15% v/v toluene compared to S12-6 and KT2440 (Figure 3). The introduction of *srp* or a combination of *slv* and *srp* enables S12-6 and KT2440 to grow in the presence of 0.3% v/v toluene. In KT2440, the introduction of both *slv* and *srp* resulted in a faster growth in the presence of 0.3% v/v toluene compared to the addition of only *srp* (Figure 3B). Interestingly, the growth of S12-6 *srp*,*slv* and S12.6 *srp* are better in comparison with S12 wildtype (Figure 3A). The observed faster growth of S12-6 *srp*,*slv* and S12.6 *srp* may be due to more efficient growth in the presence of toluene supported by a chromosomally introduced *srp* operon, compared to its original megaplasmid localization. Indeed, replication of this large megaplasmid is likely to require additional maintenance energy. To corroborate this, we complemented the megaplasmid lacking the solvent pump, pTTS12 (Tc^R^::*srpABC*) into *P. putida* S12-6 *srp* resulting in the strain *P. putida* S12-9. Indeed, *P. putida* S12-9 showed further reduced growth in the presence of 0.20 and 0.30 % toluene (Figure S2), indicating the metabolic burden of carrying the megaplasmid. We conclude that the SrpABC efflux pump can be regarded as the major contributor to solvent tolerance from pTTS12. The *slv* gene pair appears to promote tolerance of *P. putida* S12 at least under moderate solvent concentrations.

**Figure 3.**
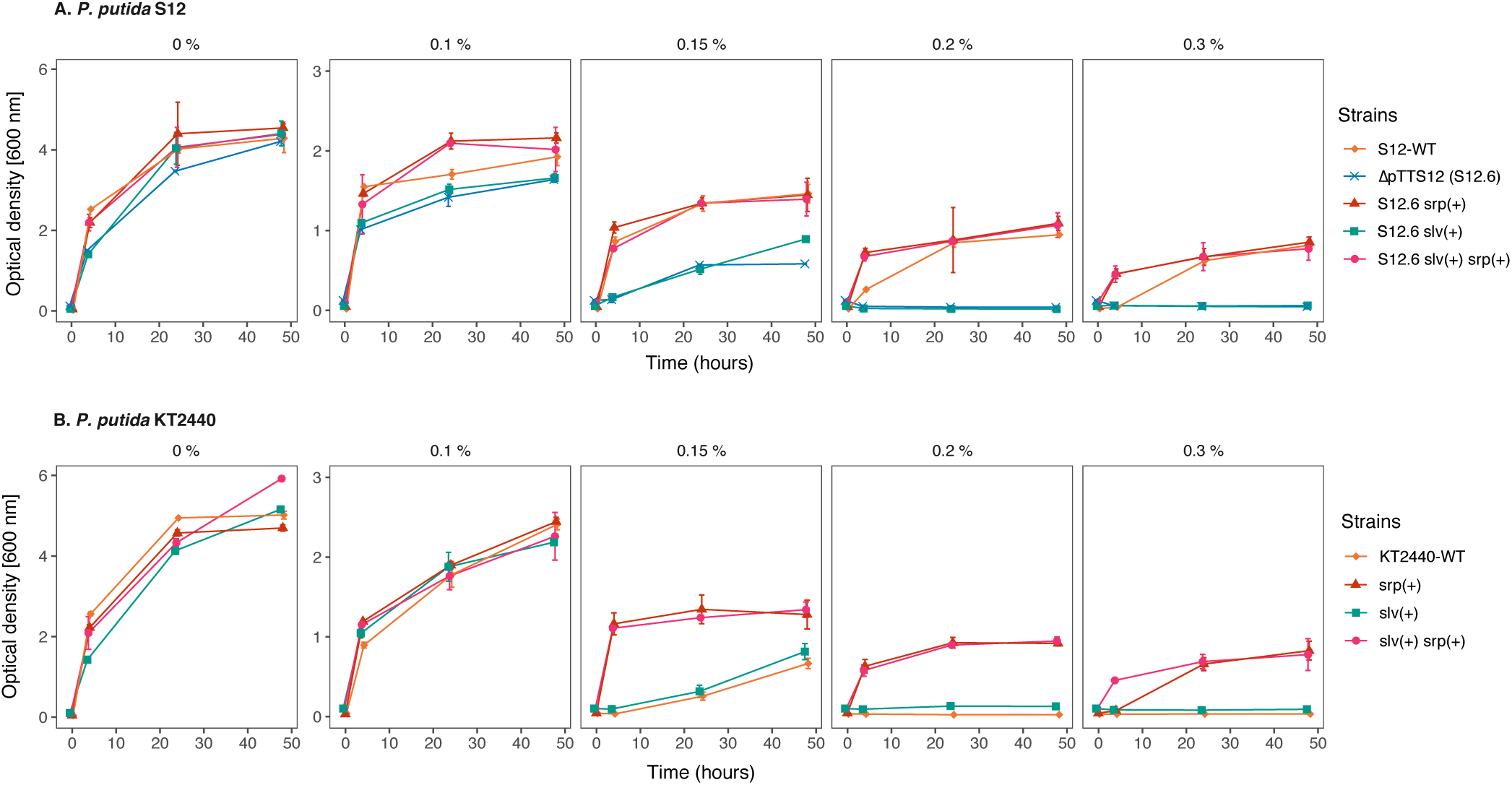
Chromosomal introduction of *srp* and *slv* gene clusters increased solvent tolerance in *P. putida* strains. Solvent tolerance analysis of the strains with chromosomal introduction of *srp* operon (*srpRSABC*), *slv* gene pair (RPPX_26255-26260) and the combination of these gene clusters into *P. putida* S12 ΔpTTS12/S12.6 (A) and wildtype *P. putida* KT2440 (B) in liquid LB with 0, 0.10, 0.15, 0.20 and 0.30 % v/v of toluene. Wildtype *P. putida* S12 was taken as a solvent tolerant control strain. This figure displays the mean of three independent replicates and error bars indicate standard deviation. The range of y-axis is different in the first panel (0–6) than the rest of the panels (0–3).

The intrinsic solvent tolerance of *E. coli* strains was observed to be clearly lower than that of *P. putida* (Figure 4). The wild type *E. coli* strains were able to withstand a maximum 0.10% v/v toluene, whereas plasmid-cured *P. putida* S12-6 and *P. putida* KT2440 were able to grow in the presence of 0.15% v/v toluene. With the introduction of *slv* and *srp* in both *E. coli* strains, solvent tolerance was increased up to 0.15% and 0.2% v/v toluene respectively (Figure 4). A combination of *slv* and *srp* also increased tolerance to 0.20% v/v toluene while showing a better growth than chromosomal introduction of just *srp*. However, none of these strains were able to grow in the presence of 0.30% v/v toluene.

**Figure 4.**
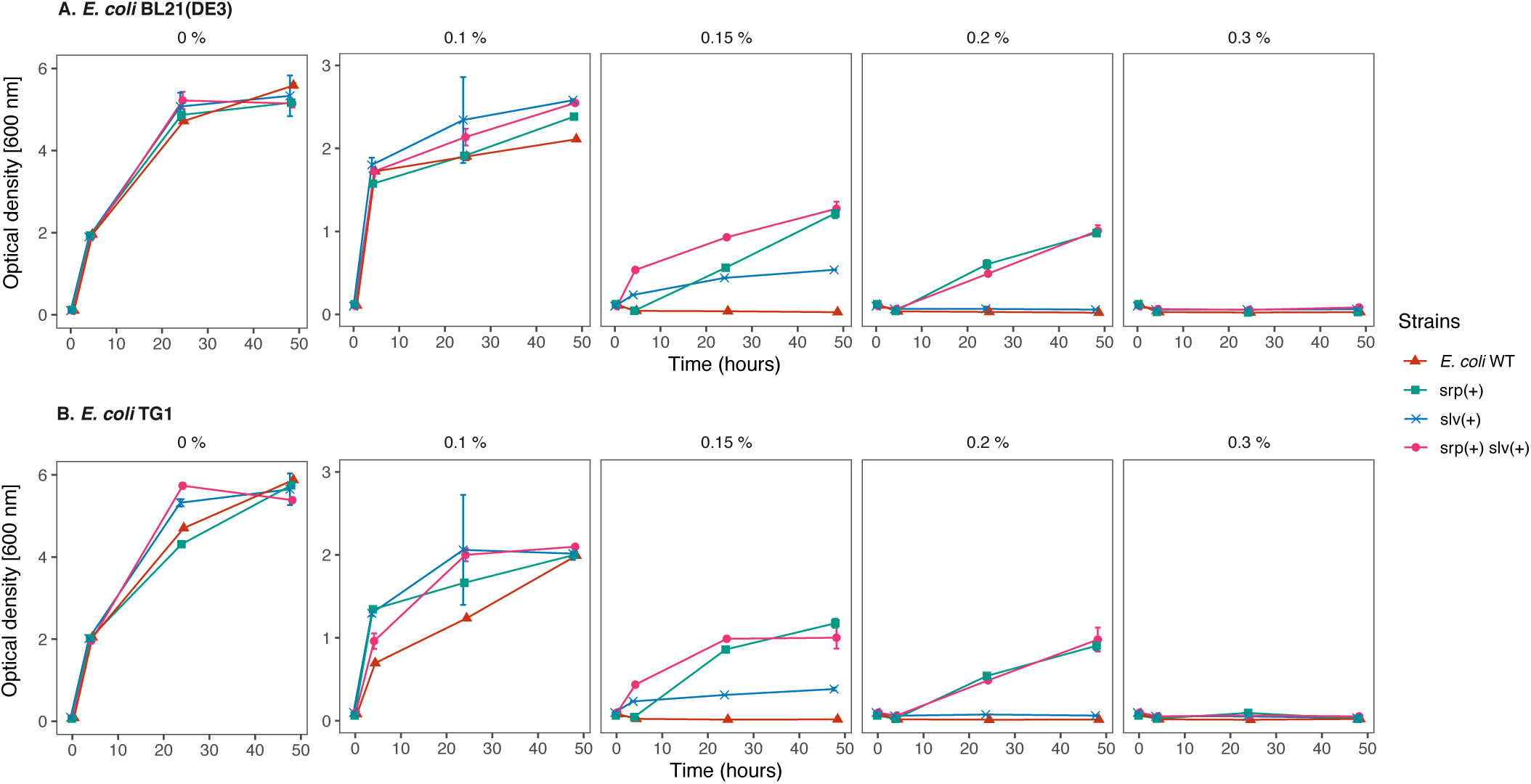
Chromosomal introduction of *srp* and *slv* gene clusters increased solvent tolerance in *E. coli* strains. Solvent tolerance analysis of the strains with chromosomal introduction of *srp* operon (*srpRSABC*), *slv* gene pair (RPPX_26255-26260) and the combination of these gene clusters into *E. coli* BL21(DE3) (A) and *E. coli* TG1 (B) in liquid LB with 0, 0.10, 0.15, 0.20 and 0.30 % v/v of toluene. This figure displays the mean of three independent replicates and error bars indicate standard deviation. The range of y-axis is different in the first panel (0–6) than the rest of the panels (0–3).

qPCR analysis of SrpABC expression (Table S1) in *P. putida* S12, *P. putida* KT2440, *E. coli* TG1, and *E. coli* BL21(DE3) confirmed that *srp*A, *srp*B, and *srp*C were expressed in basal levels in all strains. In the presence of 0.10 % toluene, the expression of *srp*A, *srp*B, and *srp*C was clearly upregulated in all strains. Thus, the lower solvent tolerance conferred by introducing SrpABC efflux pump in *E. coli* strains was not due to lower expression of the *srp* genes. Analysis of the codon adaptation index (CAI) (http://genomes.urv.es/CAIcal/) (31) showed that for both the *P. putida* and *E. coli* strains the CAI values of the srp operon are suboptimal, cleary below 0.8 to 1.0 (Table S2). Interestingly, the CAI values were higher for *E. coli* (0.664) than for *P. putida* (0.465) predicting a better protein translation efficiency of the *srp* operon in *E. coli*. Hence, reduced translation efficiency is not likely to be the cause of lower performance of *srp* operon in E. coli strains for generating solvent tolerance. Overall, our results indicate that in addition to the solvent efflux pump, *P. putida* S12 and *P. putida* KT2440 are intrinsically more robust compared to *E. coli* TG1 and *E. coli* BL21 DE3 in the presence of toluene.

### *slv* gene pair constitutes a novel toxin-antitoxin system

BLASTp analysis was initiated to further characterize RPPX_26255 and RPPX_26260. This indicated that RPPX_26260 and RPPX_26255 likely represents a novel toxin-antitoxin (TA) system. Through a database search on TADB2.0 (19, 20), we found that RPPX_26260 is a toxin of COG5654 family typically encodes a RES domain-containing protein, having a conserved Arginine (R) – Glutamine (E) – Serine (S) motive providing a putative active site and RPPX_26255 is an antitoxin of COG5642 family. Based on its involvement in solvent tolerance, we propose naming the toxin-encoding RPPX_26260 as *slvT* and the antitoxin-encoding RPPX_26255 as *slvA*.

Makarova and colleague identified putative toxin-antitoxin pairs through genome mining of reference sequences in NCBI database (32). They identified 169 pairs of the COG5654-COG5642 TA system from the reference sequences. Here, we constructed a phylogenetic tree of the COG5654-COG5642 TA system including SlvA (AJA16859.1) and SlvT (AJA16860.1) as shown in figures 5A and 6A. SlvA and SlvT cluster together with other plasmid-borne toxin-antitoxin from *Burkholderia vietnamensis* G4, *Methylibium petroleiphilum* PM1, *Rhodospirillum rubrum* ATCC 11170, *Xanthobacter autotrophicus* Py2, *Sinorhizobium meliloti* 1021, *Sinorhizobium medicae* WSM419, and *Gloeobacter violaceus* PCC7421. Multiple alignments of SlvAT against these toxin-antitoxin of COG5654-COG5642 TA system are shown in figures 5B and 6B.

**Figure 5.**
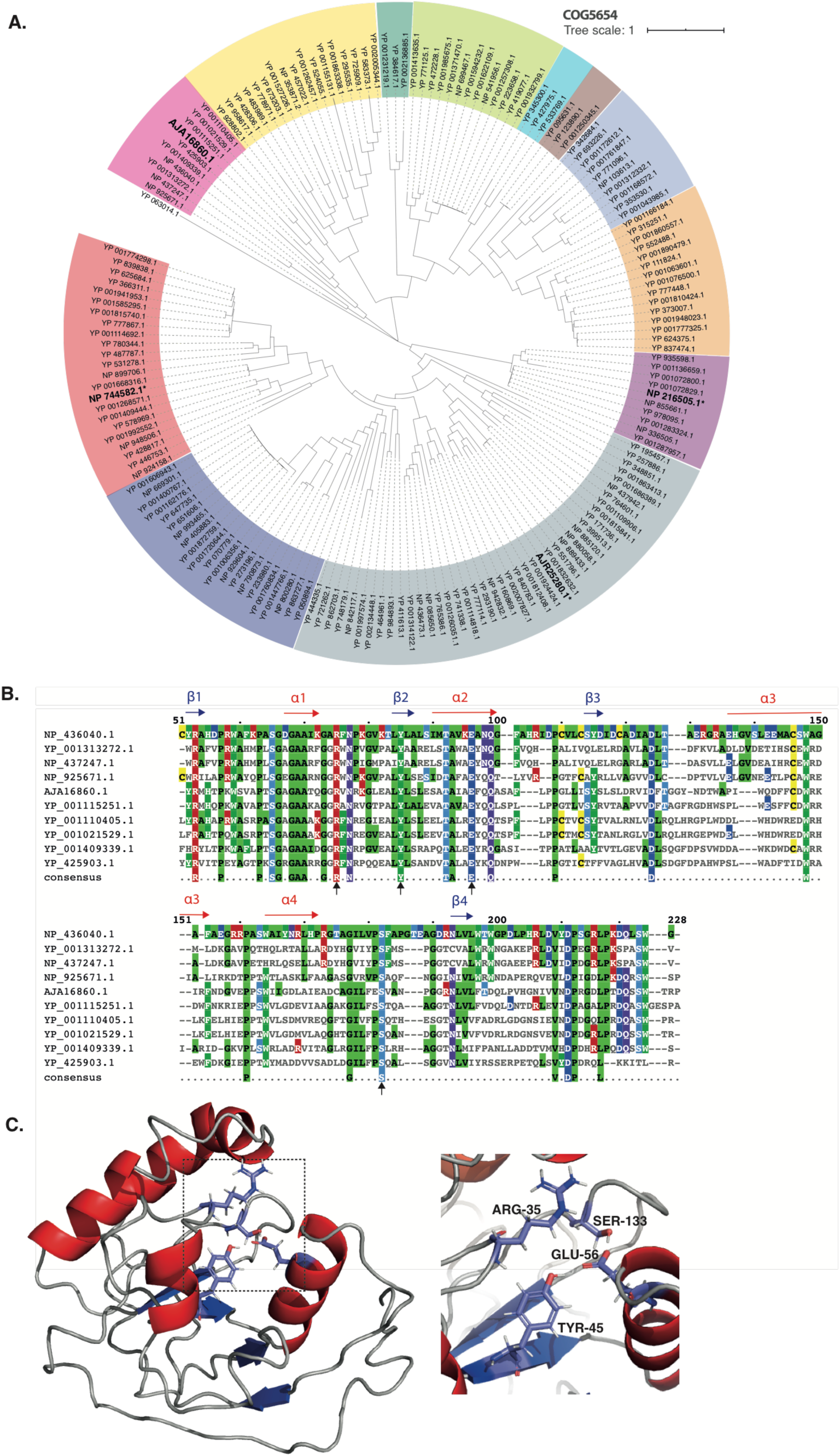
**Bioinformatics analysis of SlvT as a member of COG5654 toxin family** A. Phylogenetic tree (neighbour joining tree with 100 bootstrap) of COG5654 family toxin from reference sequences identified by Makarova and colleagues (29). Different colours correspond to the different toxin-antitoxin module clades. Asterisks (**\***) and bold text indicate the characterized toxin proteins : ParT from *Sphingobium sp.* YBL2 (AJR25280.1), PP_2434 from *P. putida* KT2440 (NP_744582.1), MbcT from *Mycobacterium tuberculosis* H37Rv (NP_216505.1), and SlvT from *P. putida* S12 (AJA16860.1). B. Multiple sequence alignment of the COG5654 toxin SlvT from *P. putida* S12 with several putative COG5654 family toxin protein which belong in the same clade. Putative active site residues are indicated by black arrows. C. Protein structure modelling of SlvT using I-TASSER server (30) which exhibits high structural similarity with MbcT from *Mycobacterium tuberculosis* H37Rv. Shown are the close up of putative active site of SlvT toxin (Arg-35, Tyr-45, Glu-56, and Ser-133).

**Figure 6.**
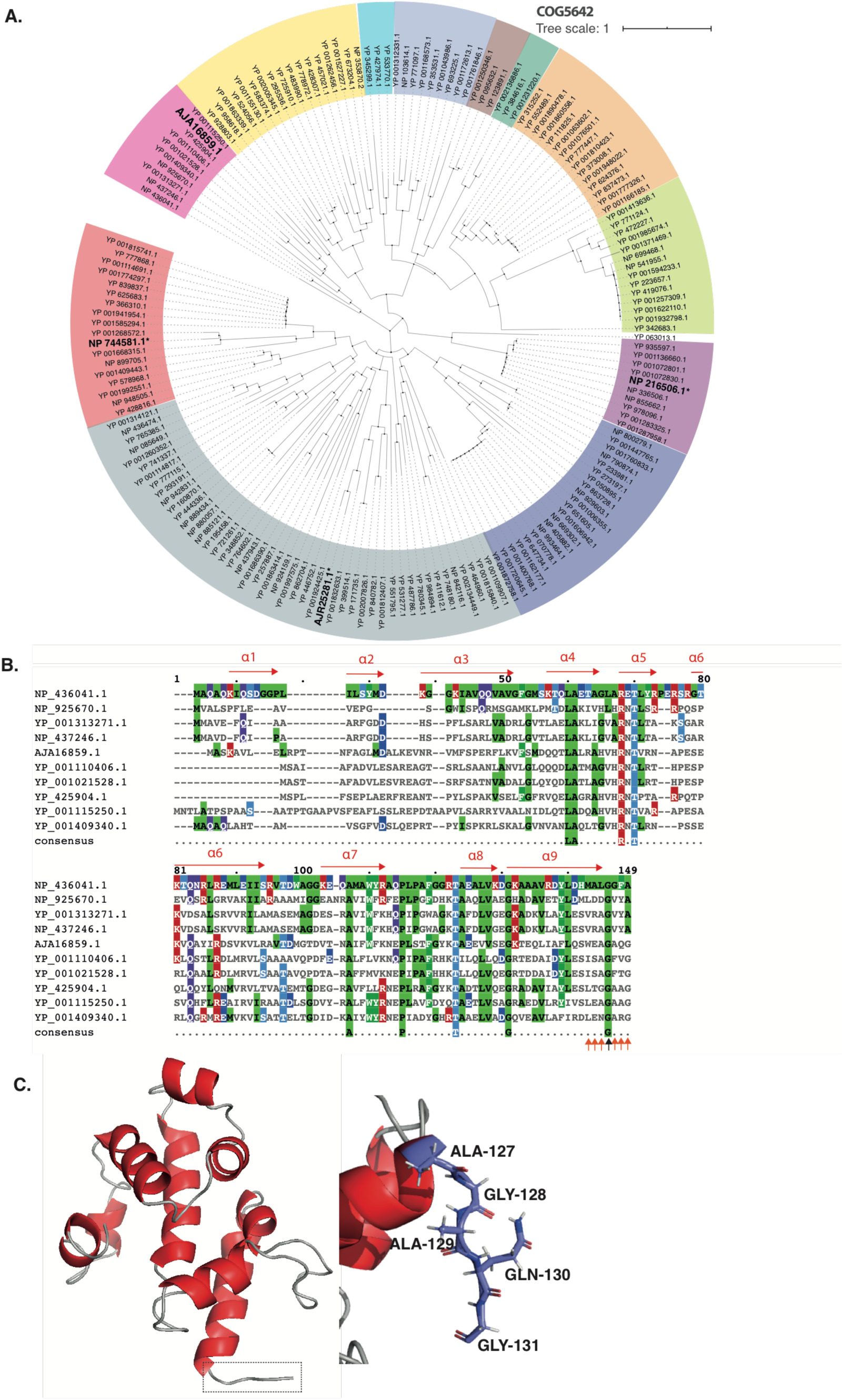
**Bioinformatics analysis of SlvA as a member of COG5642 toxin family** A. Phylogenetic tree (neighbour joining tree with 100 bootstrap) of COG5642 family toxin from reference sequences identified by Makarova and colleagues (29). Different colours correspond to the different toxin-antitoxin module clades. Asterisks (*) and bold text indicate the characterized toxin proteins : ParS from *Sphingobium sp.* YBL2 (AJR25281.1), PP_2433 from *P. putida* KT2440 (NP_744581.1), MbcA from *Mycobacterium tuberculosis* H37Rv (NP_216506.1), and SlvA from *P. putida* S12 (AJA16859.1). B. Multiple sequence alignment of the COG5654 toxin SlvA from *P. putida* S12 with several putative COG5642 family toxin protein which belong in the same clade. Putative active site residues are indicated by orange and black arrows. C. Protein structure modelling of SlvA using I-TASSER server (30) which exhibits high structural similarity with MbcA from *Mycobacterium tuberculosis* H37Rv. Shown are the close up of antitoxin putative C-terminal binding site to block SlvT toxin active site (Ala-127, Gly-128, Ala-129, Gln-130, and Gly-131).

Of the 169 TA pairs of the COG5654-COG5642 TA system, three TA pairs have recently been characterized: ParST from *Sphingobium sp.* YBL2 (AJR25281.1, AJR25280.1), PP_2433-2434 from *P. putida* KT2440 (NP_744581.1, NP_744582.1), and MbcAT from *Mycobacterium tuberculosis* H37Rv (NP_216506.1, NP216505.1) (Figure 5A and 6A, indicated by bold text and asterisks). 3D-model prediction of SlvT and SlvA protein using the I-TASSER protein prediction suite (33), indicated that SlvT and SlvA showed highest structural similarity to the MbcAT system from *Mycobacterium tuberculosis* (Figure 5C and 6C) which is reported to be expressed during stress condition (25). Amino acid conservation between SlvAT and these few characterized toxin-antitoxin pairs is relatively low, as they do not belong to the same clade (Figure 5A and 6A). However, 100% conservation is clearly observed on the putative active side residues: arginine (R) 35, tyrosine (Y) 45, and glutamine (E) 56 and only 75% consensus is shown on serine (S) 133 residue (Figure S3).

According to the model with highest TM score, SlvT is predicted to consist of four beta sheets and four alpha-helices. As such, SlvT exhibits large structural similarity with diphtheria toxin which functions as ADP-ribosyl transferase enzyme. Diphtheria toxin can degrade NAD^+^ into nicotinamide and ADP ribose (34). A similar function was recently identified for COG5654-family toxins from *P. putida* KT2440, *M. tuberculosis*, and *Sphingobium sp* (25, 26, 35).

### *slvT* toxin causes cell growth arrest by depleting cellular NAD^+^

To prove that *slvAT* presents a pair of toxin and antitoxin, *slvA* and *slvT* were cloned separately in pUK21 (lac-inducible promoter) and pBAD18 (ara-inducible promoter), respectively. The two constructs were cloned into *E. coli* BL21 (DE3). Growth of the resulting strains was monitored during conditional expression of the *slvA* and *slvT* genes (figure 6A). At the mid-log growth phase, a final concentration of 0.8% arabinose was added to the culture (*), inducing expression of *slvT*. After 2 hours of induction, growth of this strain ceased while the uninduced control culture continued to grow. Upon addition of 2 mM IPTG (**), growth of the *slvT-*induced culture was immediately restored, reaching a similar OD_600nm_ as the uninduced culture.

Bacterial cell division was further studied by flow cytometer-analyses during the expression of *slvT* and *slvA*. After approximately 6 hours of growth (indicated by grey arrow on figure 7A), samples were taken from control, arabinose, and arabinose + IPTG induced liquid culture. Cell morphology was analyzed by light microscopy and DNA content of the individual cells in the culture were measured using flow cytometer with SYBR green II staining (figure 7B). Indeed, absence of dividing cells and lower DNA content were observed during the induction of only *slvT* toxin with arabinose (figure 7B). Subsequent addition of IPTG to induce *slvA* expression was shown to restore cell division and an upshift of DNA content similar to that of control strain (figure 7B). While the expression of *slvT* was not observed to be lethal to bacterial strain, this experiment showed that the expression of *slvT* toxin stalled DNA replication and subsequently cell division. The induction of *slvA* subsequently restored bacterial DNA replication and cell division.

**Figure 7.**
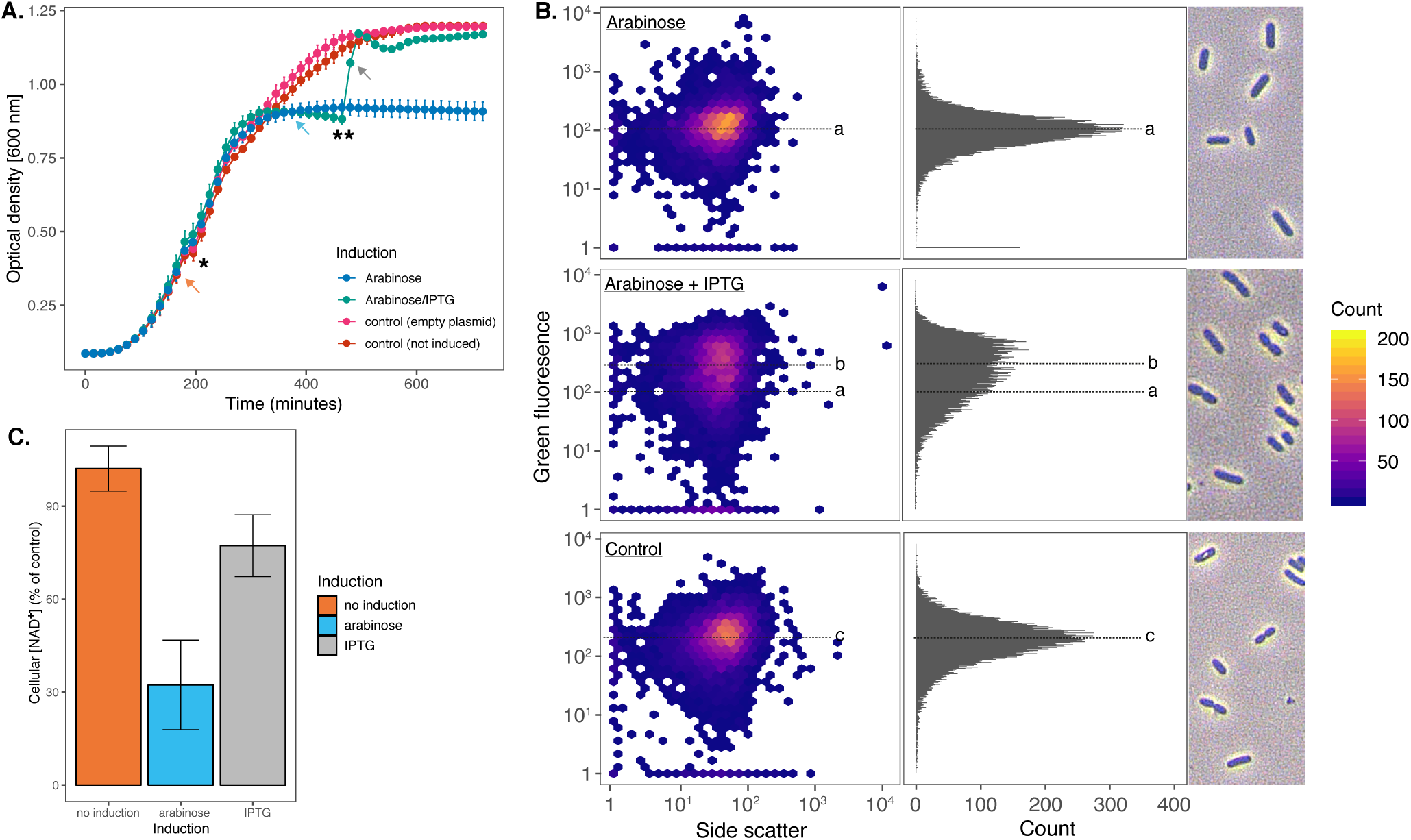
**Heterologous expression of SlvAT in *E. coli* BL21(DE3)** A. Growth curves of *E. coli* BL21(DE3) harbouring pBAD18-slvT and pUK21-slvA showing growth reduction after the induction of toxin by a total concentration of 0.8 % arabinose (*) and growth restoration after antitoxin induction by a total concentration of 2 mM IPTG (**). Samples were taken at the time points indicated by coloured arrows for cellular NAD^+^ measurement. B. Flow cytometry analysis of DNA content and cell morphology visualization on *E. coli* BL21(DE3) during *slvT* and *slvAT* expression. Median value of green fluorescence representing DNA content during *slvT* expression (118.202), *slvAT* expression (236.056), and control (208.406) are indicated by **a**, **b**, and **c** respectively. Samples were taken at the time point indicated by grey arrow on figure 6A. C. Cellular NAD^+^ measurement during the expression of toxin-antitoxin module. Induction of toxin SlvT caused a reduction in cellular NAD^+^ level to 32.32 (±14.47) % of the control strain, while the expression of SlvA restored cellular NAD^+^ level to 77.27 (±9.97) % of the control strain.

To corroborate a putative target of *slv*T, concentrations of NAD^+^ were measured during the induction experiment (figure 7C). Before the addition of arabinose to induce *slvT* (orange arrow on figure 7A), NAD^+^ was measured and compared to the strain harboring empty pUK21 and pBAD18 (figure 7B). On average, at this time point NAD^+^ level is similar between the *slvAT* bearing strain and the control strain. NAD^+^ was measured again after arabinose induction when growth of the induced strain has diminished (blue arrow on figure 7A). At this time point, the measured NAD^+^ was 32% (±14.47) of control strain. After the induction of *slvA*, NAD^+^ was immediately restored to a level of 77% (±9.97) compared to the control strain. Thus, induction of *slvT* caused depletion of NAD^+^, while induction of *slvA* immediately increased NAD^+^ level, indicating that *slvAT* is a pair of toxin-antitoxin which controls its toxicity through NAD^+^ depletion.

### *slvAT* governs megaplasmid pTTS12 stability

In addition to its role in solvent tolerance, localization of the *slvAT* pair on megaplasmid pTTS12 may have an implications for plasmid stability. pTTS12 is a very stable megaplasmid that cannot be spontaneously cured from *P. putida* S12 and cannot be removed by introducing double strand breaks (see above). We deleted *slvT* and *slvAT* from the megaplasmid to study their impact in pTTS12 stability. With the deletion of *slvT* and *slvAT*, the survival rate during treatment with mitomycin C improved significantly reaching 1.01 (± 0.17) x 10^-4^ and 1.25 (± 0.81) x 10^-4^ respectively while the wildtype S12 had a survival rate of 2.48 (± 0.58) x 10^-8^.

We determined the curing rate of pTTS12 from the surviving colonies. In wildtype S12, the curing rate was 2% (see also above) while in Δ*slvT* and Δ*slvAT* curing rate increased to 41.3% (± 4.1%) and 79.3% (± 10%) respectively, underscoring an important role for *slvAT* in megaplasmid stability. We attempted to cure megaplasmid by introducing double strand break (DSB) as previously described on *Pseudomonas taiwanensis* VLB120 (30, 36). This indeed was not possible in wildtype S12 and Δ*slvT*, however Δ*slvAT* now showed plasmid curing by DSB resulting in a curing rate of 34.3% (± 16.4%).

Since Δ*slvT* and Δ*slvAT* may compromise megaplasmid stability, we now performed megaplasmid stability tests by growing S12 and KT2440 harboring pSW-2 (negative control), pTTS12 (positive control), pTTS12 Δ*slvT*, and pTTS12 Δ*slvAT* on LB media with 10 passages (± 10 generations/passage step) as shown on figure 8. Both KT2440 and S12 easily lost the negative control plasmid pSW-2 (figure 8). Wildtype pTTS12 was not lost during this test confirming that pTTS12 is indeed a stable plasmid. Furthermore, the Δ*slvT* strains also did not show loss of megaplasmid. Interestingly, the Δ*slvAT* strains spontaneously lost the megaplasmid, confirming that the *slvAT* module is not only important to promote solvent tolerance but also determines megaplasmid stability in *P. putida* S12 and KT2440.

**Figure 8.**
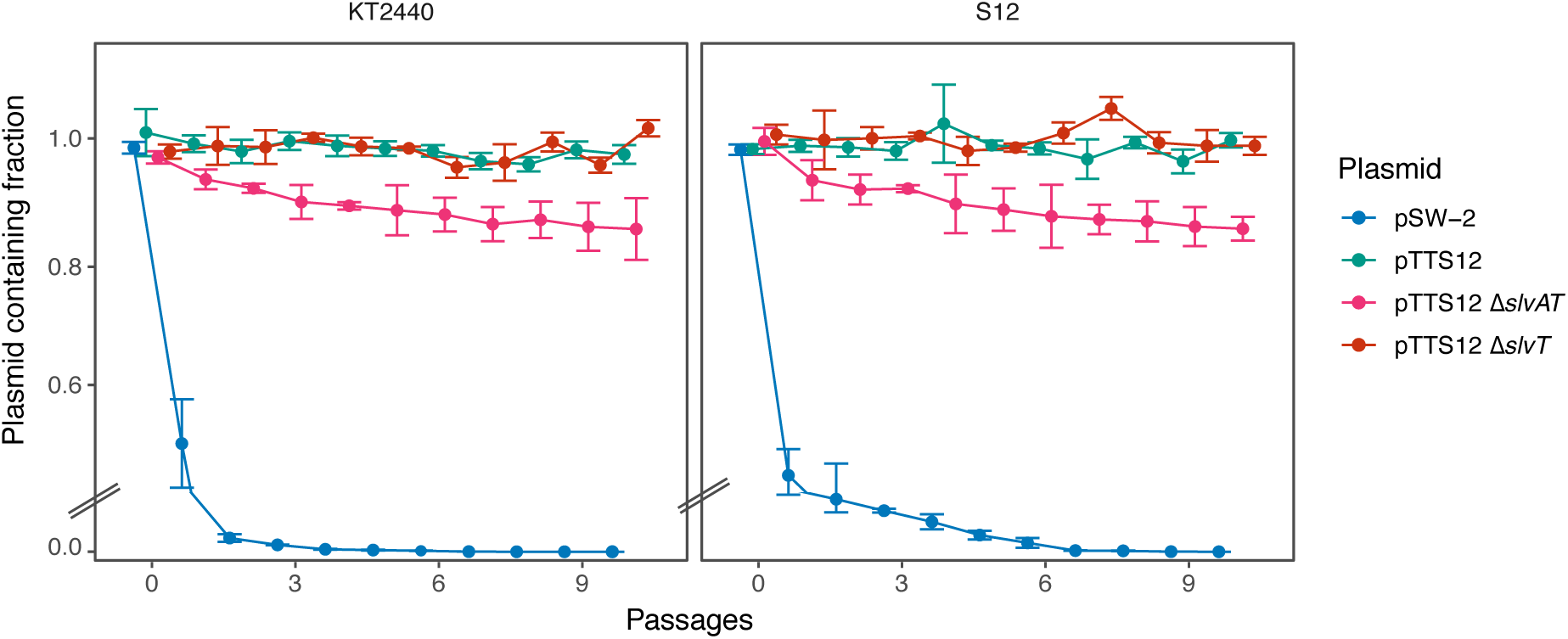
SlvAT is important for pTTS12 maintenance in *P. putida*. pTTS12 (variant with Km^R^) maintenance in *P. putida* S12 and *P. putida* KT2440 growing in LB liquid medium without antibiotic selection for 10 passages (± 10 generations per passage). pSW-2 was taken as negative control for plasmid stability in *P. putida*. This experiment was performed with three biological replicates and error bars represent standard deviation.

## Discussion

Gram-negative bacteria are regarded as preferred microbial hosts for the production of various important industrial chemicals, including biofuels and aromatic compounds. However, production of such high-value chemicals often creates an adverse effect on the cell growth due to toxicity of the produced compounds (37). Thus, in the biobased production of aromatic chemicals and biopolymers, solvent-tolerance is an essential trait for microbial hosts. In this study, removal of the megaplasmid pTTS12 from *P. putida* S12 led to the loss of the solvent-tolerant phenotype and subsequent complementation of megaplasmid pTTS12 into the plasmid-cured *P. putida* S12 restored solvent tolerance. The SrpABC efflux pump and SlvAT toxin-antitoxin module are encoded on pTTS12 and improved tolerance and survival of *P. putida* S12 to toluene exposure, making these gene clusters suitable candidates for exchange with various microbial hosts to increase tolerance towards toxic products (38).

Among all the genes encoded on the megaplasmid pTTS12, the SrpABC efflux pump appears as the major effector of solvent tolerance in *P. putida* S12 (Figure 9). A previous report applied SrpABC in whole-cell biocatalysis while optimizing the production of 1-naphtol in *E. coli* TG1 (15, 16). Implementation of the SrpABC efflux pump increased the production of 1-naphtol from E.coli, however, production was still higher using *P. putida* S12 as the production host. Here, we compared the performance of SrpABC efflux pump in several established industrial strains. SrpABC was expressed at a basal level and upregulated in the presence of 0.10 % v/v toluene in *P. putida* S12, *P. putida* KT2440, *E. coli* TG1, and *E. coli* BL21(DE3) strains. However, the *E. coli* strains clearly showed a smaller increase in toluene tolerance than the *P. putida* strains. This indicates that besides having an efficient solvent efflux pump, *P. putida* S12 and *P. putida* KT2440 are inherently more robust in the presence of toluene and, presumably, other organic solvents compared to *E. coli* TG1 and *E. coli* BL21(DE3). Detailed investigation of this intrinsic solvent tolerance of *P. putida* may further reveal the basis for this intrinsic robustness.

**Figure 9.**
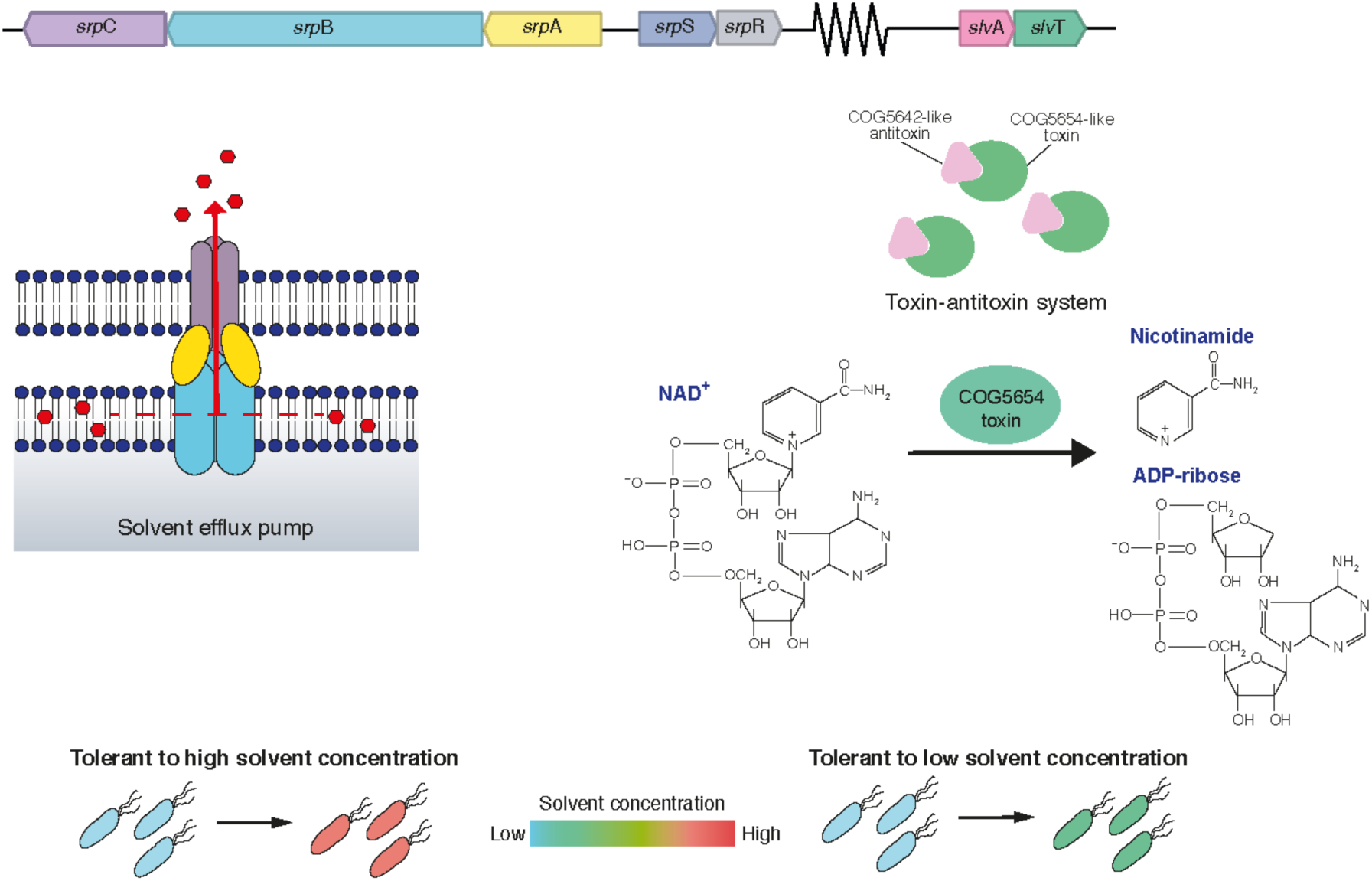
Schematic representation of the genes involved in solvent tolerance from megaplasmid pTTS12. SrpABC efflux pump is the major contributor of solvent tolerance trait from the megaplasmid pTTS12. This efflux pump is able to efficiently extrude solvents from membrane lipid bilayer. A COG5654-COG5642 family toxin-antitoxin system (SlvT and SlvA respectively) promoted the growth of *P. putida* S12 in the presence of low solvent concentration. In the absence of SlvA, SlvT causes toxicity by conferring cellular NAD^+^ depletion.

We recently identified two genes upregulated in transcriptome analysis of toluene-shocked *P. putida*, RPPX_26255 and RPPX_26260, putatively playing a role in solvent tolerance (11). Here, we confirmed this finding and demonstrated that these genes together form a novel toxin-antitoxin module (Figure 7). Sequence comparison with other known toxin-antitoxin gene pairs in the toxin-antitoxin database TADB2.0, revealed that RPPX_26255 (*slvA*) contains a DUF2834 domain characteristic for COG5642-family antitoxin, while RPPX_26260 (*slvT*) carries a conserved RES domain like other COG5654-family toxins (19, 20). In accordance to this result, structural similarity of SlvT and SlvA with other toxin-antitoxin pairs of the COG5654-COG5642 family was confirmed through 3D-structure prediction with the protein structure and function prediction tool I-TASSER (33).

Toxin-antitoxin systems are known to be important in antibiotic persistent strains as a trigger to enter and exit the dormant state, causing the cell to become unaffected by the antibiotic (39). Among *Pseudomonas* species, several toxin-antitoxin systems are reported to be involved in survival strategies, such as stress response, biofilm formation, and antimicrobial persistence (27, 40–42). In this paper, we show that the novel toxin-antitoxin system represented by SlvAT improves solvent tolerance and is important for megaplasmid pTTS12 stability. SlvT exerts toxicity by degradation of NAD^+^, like other toxins of the COG5654-family, and expression of antitoxin SlvA immediately restored NAD^+^ levels. Depletion of NAD^+^ interfered with DNA replication and caused arrest of cell division similar to another recently described COG5654-COG5642 family toxin-antitoxin pair (27).

Megaplasmids, such as pTTS12, may cause a metabolic burden for the strains that harbor them, and such plasmid can be a source of genetic instability (43). We show that pTTS12 indeed imposed a metabolic burden in the presence of organic solvent. In addition, we demonstrated the importance of the SlvAT toxin-antitoxin module for the stabilization and maintenance of the megaplasmid which contains several gene clusters responsible for efficient stress tolerance phenotypes. Future research is required to reveal details of the control mechanisms operating in balanced in vivo.

In summary, our experiments confirmed that the SrpABC efflux pump is the major contributor of solvent tolerance on the megaplasmid pTTS12. In addition, the megaplasmid carries a novel toxin-antitoxin system SlvAT (RPPX_26255 and RPPX_26260) which promotes solvent tolerance in *P. putida* S12 and is important to maintain genetic stability of pTTS12. Chromosomal introduction of the *srpABC* operon genes in combination with *slvAT* confers a clear solvent tolerance phenotype in other industrial strains previously lacking this phenotype such as *P. putida* KT2440, *E. coli* TG1, and *E. coli* BL21(DE3).

## Materials and Methods

### Strains and culture conditions

Strains and plasmids used in this paper are listed in Table S1. All *P. putida* strains were grown in Lysogeny Broth (LB) on 30 °C with 200 rpm shaking. *E. coli* strains were cultivated in LB on 37 °C with 250 rpm. For solid cultivation, 1.5 % (w/v) agar was added to LB. M9 minimal medium was supplemented with 2 mg MgSO4 and 0.2 % of citrate as sole carbon source (44). Toluene atmosphere growth was evaluated on solid LB media in a glass plate incubated in an exicator with toluene supplied through the gas phase at 30 °C. Solvent tolerance analysis was performed by growing *P. putida* S12 strains in LB starting from OD600 0.1 in Boston bottles with Mininert bottle caps. When required, gentamycin (25 mg L^-1^), ampicillin (100 mg L^-1^), kanamycin (50 mg L^-1^), indole (100 g L^-1^), potassium tellurite (6.75-200 mg L^-1^), arabinose (0.8% m/v), and IPTG (2 mM) were added to the media.

### DNA and RNA methods

All PCRs were performed using Phusion polymerase (Thermo Fisher) according to the manufacturer’s manual. Primers used in this paper (Table S3) were procured from Sigma-Aldrich. PCR products were checked by gel electrophoresis on 1 % (w/v) TBE agarose containing 5 µg mL^-1^ ethidium bromide (110V, 0.5x TBE running buffer). For RT-qPCR analysis, RNA was extracted using TRIzol reagent (Invitrogen) according to the manufacturer’s manual. The obtained RNA samples were immediately reverse transcribed using iScript^TM^ cDNA synthesis kit (BioRad) and cDNA may be stored at -20 °C prior to qPCR analysis. qPCR was performed using iTaq^TM^ Universal SYBR Green Supermix (BioRad) on CFX96 Touch^TM^ Real-Time PCR Detection System (BioRad). The genome sequence of *P. putida* S12 ΔpTTS12 was analysed using Illumina HiSeq (GenomeScan BV, The Netherlands) and assembled according to the existing complete genome sequence (Accession no. CP009974 and CP009975) (12). These sequence data have been submitted to the DDBJ/EMBL/GenBank databases under accession number ….

### Curing and complementation of megaplasmid pTTS12 from *P. putida* S12

*P. putida* S12 was grown in LB to reach early exponential phase (± 3 hours or OD_600nm_ 0.4-0.6). Subsequently, mitomycin C was added to the liquid LB culture to a final concentration range of 10, 20, 30, 40, or 50 μg/ml. These cultures were grown for 24 hours and plated on M9 minimal media supplemented with indole to select for the absence of megaplasmid. Loss of megaplasmid was confirmed by loss of other phenotypes connected with the megaplasmid such as MIC reduction of potassium tellurite and solvent sensitivity under toluene atmosphere, as well as through genomic DNA sequencing. Complementation of megaplasmid pTTS12 was performed using bi-parental mating between *P. putida* S12-1 (pTTS12 Km^R^) and plasmid-cured strains *P. putida* S12 ΔpTTS12 (Gm^R^ :: Tn7) and followed by selection on LB agar supplemented with Kanamycin and Gentamicin.

### Plasmid cloning

Deletion of *srpABC*, *slvT*, and *slvAT* genes was performed using homologous recombination between free-ended DNA sequences that are generated by cleavage on unique I-SceI sites (36). Two homologous recombination sites were chosen downstream (TS-1) and upstream (TS-2) of the target genes. TS-1 and TS-2 fragments were obtained by performing PCR using primers listed in Table S1. Constructs were verified by DNA sequencing. Mating was performed as described by Wynands and colleagues (30). Deletion of *srpABC*, *slvT*, and *slvAT* was verified by PCR and Sanger sequending (Macrogen B.V., Amsterdam).

Introduction of the complete *srp* operon (*srpRSABC*) and *slvAT* was accomplished using the mini-Tn7 delivery vector backbone of pBG35 developed by Zobel and colleagues (45). The DNA fragments were obtained by PCR using primer pairs listed on Table S3 and ligated into pBG35 plasmid at PacI and XbaI restriction site. This construct generated a Tn7 transposon segment in pBG35 containing gentamycin resistance marker and *srp* operon with Tn7 recognition sites flanking on 5’ and 3’ sides of the segment. Restriction analysis followed by DNA sequencing (Macrogen, The Netherlands) were performed to confirm the correct pBG-srp, pBG-slv, and pBG-srp-slv construct. The resulting construct was cloned in *E. coli* WM3064 and introduced into *P. putida* or *E. coli* strains with the help of *E. coli* WM3064 pTnS-1. Integration of construct into Tn7 transposon segment was confirmed by gentamicin resistance, PCR, and the ability of the resulting transformants to withstand and grow under toluene atmosphere.

### Toxin-antitoxin assay

Bacterial growth during toxin-antitoxin assay was obeserved in LB media supplemented with 100 mg L^-1^ ampicillin and 50 mg L^-1^ kanamycin. Starting cultures were innoculated from 1:100 dilution of overnight culture (OD600 ± 0.1) into a microtiter plate (96 well) and bacterial growth was measured using Tecan Spark^TM^ 10M. To induce toxin and antitoxin, a total concentration of 0.8% m/v arabinose and 2 mM IPTG were added to the culture respectively. Cell morphology was observed using light microscope (Zeiss Axiolab 5) at 100x magnification. A final concentration of 2.5x SYBR Green I (10000x stock, New England Biolabs) was applied to visualize DNA, followed by two times washing with 1x phosphate buffer saline (PBS), and analyzed using a Guava® easyCyte Single Sample Flow Cytometer (Millipore). At indicated time points, NAD^+^ levels were measured using NAD/NADH-Glo^TM^ assay kit (Promega) according to the manufacturer’s manual. RPPX_26255 and RPPX_26260 was modelled using I-TASSER server (33) and visualized using PyMol (version 2.3.1). Phylogenetic trees of toxin-antitoxin module derived from COG5654-COG5642 family were constructed using MEGA (version 10.0.5) as a maximum likelihood tree with 100 bootstrap and visualized using iTOL webserver (https://itol.embl.de) (46).

**Table 1.** Strains and plasmids used in this paper

| Strain | Characteristics | References |
| --- | --- | --- |
| *P. putida* S12 | Wild type *P. putida* S12 (ATCC 700801), harboring megaplasmid pTTS12 | (3) |
| *P. putida* S12-1 | *P. putida* S12, harboring megaplasmid pTTS12 with Km<sup>R</sup> marker | This paper |
| *P. putida* S12-6/ S12-10/ S12-22 | ΔpTTS12 | This paper |
| *P. putida* S12-9 | ΔpTTS12, Gm<sup>R</sup> *srpRSABC*::Tn7, complemented with megaplasmid pTTS12 (Tc<sup>R</sup>::*srpABC*) | This paper |
| *P. putida* S12-C | *P. putida* ΔpTTS12 (S12-6/ S12-10/ S12-22), complemented with megaplasmid pTTS12 | This paper |
| *P. putida* KT2440 | Derived from wildtype *P. putida* mt-2, ΔpWW0 | (47) |
| *E. coli* HB101 | *recA pro leu hsdR* Sm<sup>R</sup> | (48) |
| *E. coli* BL21(DE3) | *E. coli* B, F<sup>-</sup> *ompT gal dcm lon hsdS<sub>B</sub>(r<sub>B</sub><sup>-</sup>m<sub>B</sub><sup>-</sup>)* λ(DE3 [*lacI lacUV5-T7p07 ind1 sam7 nin5*]) [*malB*<sup>+</sup>]<sub>K-12</sub>(λ<sup>S</sup>) | (49) |
| *E. coli* DH5α λpir | *sup* E44, Δ*lacU*169 (Φ*lacZ*ΔM15), *recA1, endA1, hsdR17, thi-1, gyrA96, relA1*, λpir phage lysogen | (50) |
| *E. coli* TG1 | *E. coli* K-12, *glnV44 thi-1* Δ(*lac-proAB*) Δ(*mcrB-hsdSM*)5(*r<sub>K</sub><sup>-</sup>m<sub>K</sub><sup>-</sup>*) F' [*traD36 proAB<sup>+</sup> lacI<sup>q</sup> lacZ*ΔM15] | Lucigen |
| *E. coli* WM3064 | *thrB1004 pro thi rpsL hsdS lacZ*ΔM15 RP4-1360 Δ(*araBAD*)567 Δ*dapA*1341::[erm pir] | William Metcalf |
| **Plasmid** |  |  |
| pRK2013 | RK2-Tra<sup>+</sup>, RK2-Mob<sup>+</sup>, Km<sup>R</sup>, *ori* ColE1 | (51) |
| pEMG | Km<sup>R</sup>, Ap<sup>R</sup>, *ori* R6K, *lacZα* MCS flanked by two I-SceI sites | (36) |
| pEMG-Δ*srpABC* | pEMG plasmid for constructing *P. putida* S12 Δ*srpABC* | This paper |
| pEMG-Δ*slvAT* | pEMG plasmid for constructing *P. putida* S12 Δ*slvAT* | This paper |
| pEMG-Δ*slvT* | pEMG plasmid for constructing *P. putida* S12 Δ*slvT* | This paper |
| pSW-2 | Gm<sup>R</sup>, *ori* RK2, *xyIS*, Pm → I-sceI | (36) |
| pBG35 | Km<sup>R</sup>, Gm<sup>R</sup>, *ori* R6K, pBG-derived | (45) |
| pBG-srp | Km<sup>R</sup>, Gm<sup>R</sup>, *ori* R6K, pBG-derived, contains *srp* operon (RPPX_27995-27965) | This paper |
| pBG-slv | Km<sup>R</sup>, Gm<sup>R</sup>, *ori* R6K, pBG-derived, contains *slv* gene pair (RPPX_26255-26260) | This paper |
| pBG-srp-slv | Km<sup>R</sup>, Gm<sup>R</sup>, *ori* R6K, pBG-derived, contains *slv* gene pair (RPPX_26255-26260) and *srp* operon (RPPX_27995-27965) | This paper |
| pBAD18-slvT | Ap<sup>R</sup>, *ara* operon, contains *slvT* (RPPX_26260) | This paper |
| pUK21-slvA | Km<sup>R</sup>, lac operon, contains *slvA* (RPPX_26255) | This paper |
| pTnS-1 | Ap<sup>R</sup>, *ori* R6K, TnSABC+D operon | (52) |

## Acknowledgement

H. Kusumawardhani was supported by the Indonesia Endowment Fund for Education (LPDP) as scholarship provider from the Ministry of Finance, Indonesia. R. Hosseini was funded by the Dutch National Organization for Scientific Research NWO, through the ERAnet-Industrial Biotechnology program, project ‘Pseudomonas 2.0’.

## List of supplementary materials

**Table S1. Expression of *srpABC* genes in *P. putida* and *E. coli* strains in basal level and in the presence of 10 mM toluene with *gyrB* and *rpoB* as reference genes**

**Table S2. Codon adaptation index of *srp* operon in *E. coli* and *P. putida* reference strains**

**Table S3. Primers used in this paper**

**Figure S1. Removal and complementation of the megaplasmid pTTS12 from *P. putida* S12.**

A. The loss of the megaplasmid band in megaplasmid-cured *P. putida* S12 proven by electrophoresis of agarose embedded genomic DNA. Megaplasmid band (orange arrow) was visible in the positive control *P. putida* S12 and absent in negative control *P. putida* KT2440 and Mitomycin C treated strains (strain S12-6, S12-10, and S12-22). Blue arrow indicates bacterial chromosome.

B. Activity of styrene monooxygenase (SMO) and styrene oxide isomerase (SOI) for indigo formation from indole in *P. putida* strains. Enzyme activity was lost in the megaplasmid-cured strains S12 ΔpTTS12 (white colonies) and restored with the complementation of megaplasmid in the strains S12-C (blue colonies). Indole (100 mg L^-1^) was supplemented in M9 minimum media.

C. K_2_TeO_3_ resistance of *P. putida* strains on lysogeny broth (LB) agar. Tellurite resistance was reduced in the megaplasmid-cured strains S12 ΔpTTS12 (MIC 50 mg L^-1^) and restored with the complementation of megaplasmid in the strains S12-C (MIC 200 mg L^-1^).

D. Solvent tolerance analysis was performed on *P. putida* S12, *P. putida* S12 ΔpTTS12, and *P. putida* S12-C growing in liquid LB media with 0, 0.10, 0.15, 0.20 and 0.30 % v/v toluene. The removal of the megaplasmid pTTS12 clearly caused a significant reduction in the solvent tolerance of *P. putida* S12 ΔpTTS12. Complementation of pTTS12 restores the solvent tolerance trait in *P. putida* S12-C. This figure displays the mean of three independent replicates and error bars indicate standard deviation. The range of y-axis is different in the first panel (0–6) than the rest of the panels (0 - 2.5).

**Figure S2. Metabolic burden of megaplasmid pTTS12 during growth in the presence of organic solvent.**

Solvent tolerance was compared between *P. putida* S12, *P. putida* S12-6.1 (S12-6 srp::attn7), and *P. putida* S12-9 (S12-6 srp::attn7, pTTS12 tet::srp) in liquid LB media with 0, 0.10, 0.15, and 0.20 % v/v toluene. This figure displays the mean of three independent replicates and error bars indicate standard deviation. The range of y-axis is different in the first panel (0–6) than the rest of the panels (0 - 2.5).

**Figure S3. Multiple alignment of SlvT and SlvA with characterized toxin-antitoxin of COG5654-COG5642 family**

Sequence similarity of the COG5654 toxin SlvT (A) from P. putida S12 and COG5642 antitoxin SlvA (B) with several characterized COG5654-COG5642 family toxin-antitoxin protein. Putative active site residues showed >70% similarities and are indicated by red arrows.

